# Pharmacologic profiling reveals lapatinib as a novel antiviral against SARS-CoV-2 in vitro

**DOI:** 10.1101/2020.11.25.398859

**Authors:** M. H. Raymonda, J. H. Ciesla, M. Monaghan, J. Leach, G. Asantewaa, L.A. Smorodintsev-Schiller, M. M. Lutz, X. L. Schafer, T. Takimoto, S. Dewhurst, J. Munger, I. S. Harris

## Abstract

The emergence of SARS-CoV-2 virus has resulted in a worldwide pandemic, but an effective antiviral therapy has yet to be discovered. To improve treatment options, we conducted a high-throughput drug repurposing screen to uncover compounds that block the viral activity of SARS-CoV-2. A minimally pathogenic human betacoronavirus (OC43) was used to infect physiologically-relevant human pulmonary fibroblasts (MRC5) to facilitate rapid antiviral discovery in a preclinical model. Comprehensive profiling was conducted on more than 600 compounds, with each compound arrayed at 10 dose points (ranging from 20 μM to 1 nM). Our screening revealed several FDA-approved agents that act as novel antivirals that block both OC43 and SARS-CoV-2 viral replication, including lapatinib, doramapimod, and 17-AAG. Importantly, lapatinib inhibited SARS-CoV-2 replication by over 50,000-fold without any toxicity and at doses readily achievable in human tissues. Further, both lapatinib and doramapimod could be combined with remdesivir to dramatically improve antiviral activity in cells. These findings reveal novel treatment options for people infected with SARS-CoV-2 that can be readily implemented during the pandemic.

## INTRODUCTION

In 2020, from February to late November, the COVID-19/SARS-CoV-2 pandemic will have killed over 256,000 people in the United States (CDC data tracker) and over 1,368,000 people globally (WHO Operational Update). Currently, effective treatments to inhibit SARS-CoV-2 morbidity and mortality are not available, and as such, the pandemic is predicted to continue to take a devastating toll on both human health and economic activity.

Since the pandemic began, there has been a strong interest in screening existing compound libraries for their ability to inhibit SARS-CoV-2 replication. Traditionally, high throughput antiviral screens test compounds in a host cell line that can be efficiently infected by the virus and is capable of supporting high titer viral replication (Bojkova et al., 2020; Gordon et al., 2020). Many times, this results in the host cell type being derived from a non-physiological source, e.g., from a different species (monkey), tissue (kidney), or genetic background (tumors) than what is important for human infection and the associated pathology. This has been the case for many SARS-CoV-2 screens, which have relied on monkey kidney cells, e.g., Vero cells, to identify potential SARS-CoV-2 antiviral agents (Bojkova et al., 2020; Gordon et al., 2020). Further, recent findings suggest these models are insufficient to identify effective SARS-CoV-2 antivirals in human cells (Dittmar, Jun 19, 2020). In addition, many screening efforts have focused on screening libraries that contain compounds at a single high concentration, e.g., 5 or 10 μM (Riva et al., 2020). While this is useful for drug discovery efforts using biochemical assays, these libraries are poorly suited for more complicated screens with live cells, as high drug concentrations can potentially reduce cell viability and induce off-target effects. Screening at only a single high dose, therefore, may completely miss many compounds that may be effective in limiting viral replication at lower concentrations.

To address these issues, we developed an anti-coronaviral screening platform in which physiologically-relevant non-transformed human pulmonary fibroblasts infected with coronavirus were tested against compounds across a wide range of doses (20 μM to 1 nM). This enabled us to identify concentrations of compounds that inhibit virally-induced killing while also maintaining cell viability. Similar drug screening approaches have successfully identified selective vulnerabilities in cancer cells (Harris et al., 2019; Nicholson et al., 2019; Shu et al., 2020). To facilitate rapid screening, we tested the minimally pathogenic human coronavirus, OC43, which belongs to the same betacoronavirus genus as SARS-CoV-2. We then examined the ability of the compound hits from our high-throughput screen to inhibit SAR-CoV-2 infection. This approach selects for antiviral compounds that may exhibit broad activity towards coronaviruses. We have identified and validated several compounds that inhibit the replication of OC43 and SARS-CoV-2. Of particular note, we find that lapatinib, an FDA-approved drug that exhibits a good toxicity profile in humans (Spector et al., 2015), inhibits SARS-CoV-2 RNA replication by over 50,000-fold.

## RESULTS

### High-throughput screening reveals compounds with novel activity against coronaviruses

Since a BSL3 high-throughput screening facility was not readily available, compounds were screened against the virus OC43, a minimally pathogenic human betacoronavirus of the same coronavirus genus as SARS-CoV-2. We tested whether OC43 would replicate in MRC5-hT cells (herein referred to as MRC5 cells), which are human pulmonary fibroblasts (immortalized with human TERT) that have previously been shown to be an effective model of viral infection into non-transformed cells (Rodriguez-Sanchez et al., 2019). The OC43 virus rapidly replicated in MRC5 cells as indicated by increased viral RNA and protein abundance in cells upon infection (Fig. S1A and S1B). Viral replication causes cell death, known as the cytopathic effect (CPE). As a readout of the ability of a compound to block viral replication, inhibition of CPE was quantified using an image-based readout. OC43 induced substantial CPE in MRC5 cells (Fig. 1A), and conditions were optimized to generate a robust Z’ factor for the screen (Zhang et al., 1999) (Z’ = 0.66) (Fig. S2). To comprehensively understand the pharmacologic profile for each compound, a new platform was developed, termed the Multifunctional Approach to Pharmacologic Screening (MAPS) (Fig. 1B). Here, each drug in our compound library was arrayed across 10 dose points, ranging from 20 μM to 1 nM. These multiple concentrations provided a broad picture of each drugs’ ability to block virus-induced CPE. For our screen, we repurposed a drug library that focused on cancer and metabolic targets, which previously had been used to examined drug sensitivities in cancer cells (Harris et al., 2019; Nicholson et al., 2019; Shu et al., 2020). The maximal inhibition of viral killing in cells (at any dose) was used to identify compounds as potential hits (Fig. 1C). Drugs spanning a wide range of target proteins were identified as antivirals (Fig. 1D). Three of the top-scoring compounds were doramapimod (BIRB 796), lapatinib, and 17-AAG. Doramapimod is a pan-inhibitor of p38 MAPKs (Pargellis et al., 2002), lapatinib is a dual inhibitor of EGFR/HER2 (Burris et al., 2005), and 17-AAG blocks HSP90 activity (Kamal et al., 2003). Upon validation using a larger number of drug concentrations, the hit compounds demonstrated robust inhibition of CPE (Fig. 1E). Interestingly, some compounds, such as 17-AAG, demonstrated that screening at μM concentrations precluded the ability to detect any rescue from viral killing due to inherent cellular toxicity. These findings illustrate the power of the MAPS platform, which identified hits that would have been most likely missed if the screen was performed at a single high drug concentration.

**Figure 1.**
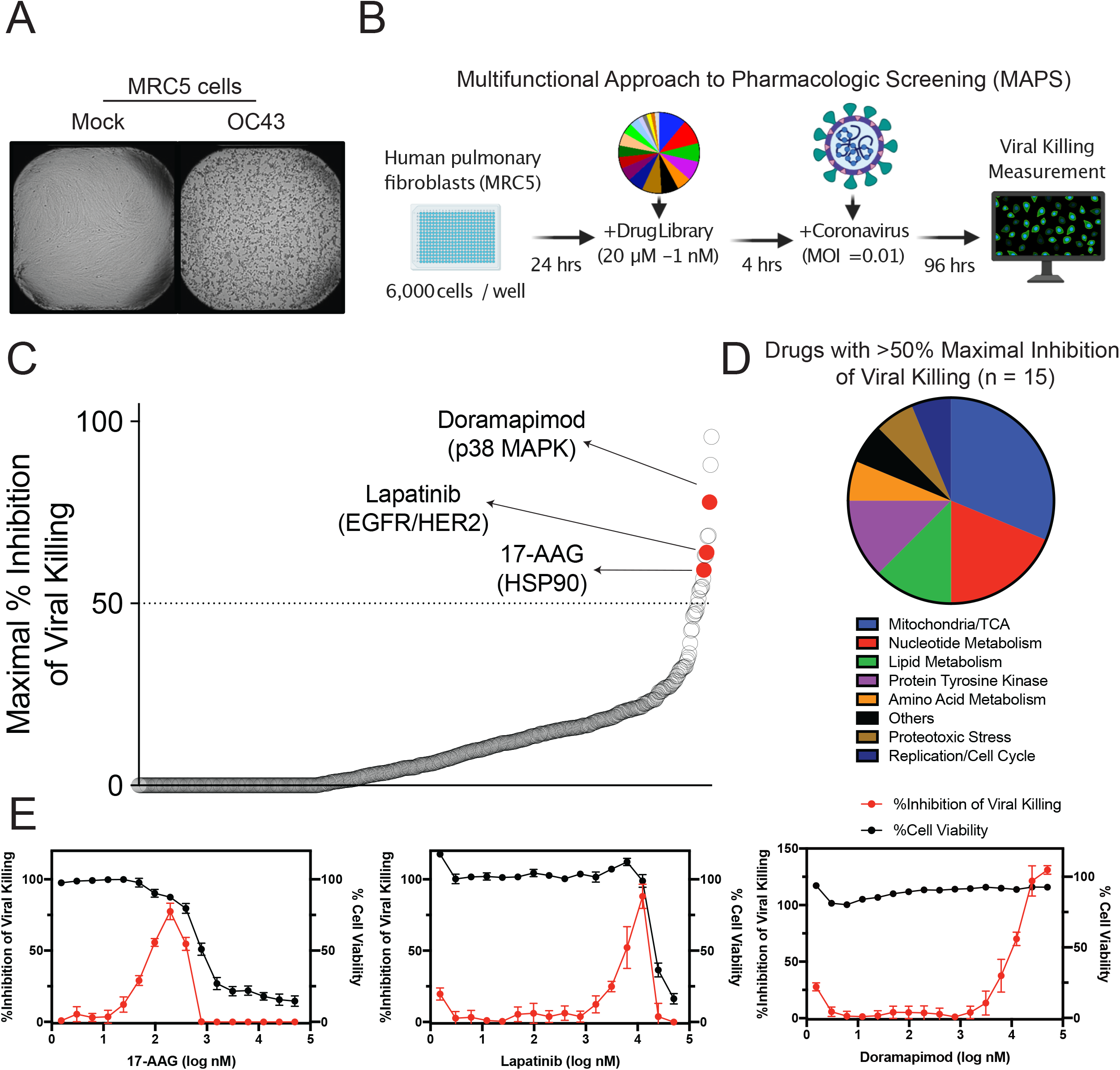
Inhibitors of viral-induced killing identified using high-throughput screening. **(A)** MRC5 cells were infected with OC43 at an MOI of 0.05 TCID50/mL. Representative images of MRC5 96 hours post infection. **(B)** Schematic of the Multifunctional Approach to Pharmacologic Screening platform used to detect compounds that inhibit coronavirus viral killing. Fibroblasts are seeded on a 384-well plate, incubated for 24 hours, treated with the drug library and then infected with coronavirus. The ability of the compounds to inhibit viral killing is measured 96 hours after infection by imaging the wells and counting nuclei. **(C)** Ranking of library compounds according to the maximal percent inhibition of viral killing across all concentrations. **(D)** Pathways associated with the 15 compounds observed to inhibit viral killing by at least 50%. **(E)** Cell viability (black) and % inhibition of viral killing (red) plotted against drug concentration for top hits 17-AAG, lapatanib and doramapimod. See also **Figure S1 and S2**.

### Hit compounds block coronavirus replication and synergize with remdesivir

Hit compounds were identified through high-throughput MAPS screening using reduction of CPE as a readout for viral inhibition. The antiviral activity of so-identified compounds was further validated by measuring the production of viral RNA and the cells’ ability to produce infectious virus (TCID50). A dose-dependent reduction in viral RNA accumulation and TCID50 was observed across all drugs (Fig. 2A-2B). Interestingly, 17-AAG was able to dramatically block viral RNA accumulation at doses almost 10-fold less than remdesivir, an antiviral with potent activity against SARS-CoV-2 in vitro. Next, we investigated whether our hit compounds could be combined with remdesivir to synergistically block CPE. While remdesivir can block SARS-CoV-2 in vitro, recent clinical trials have shown it not to be effective in patients (Pan et al., 2020). One potential limitation of remdesivir treatment is that its IC50 against SARS-CoV-2 is in the μM range, and plasma concentrations in COVID-19 infected patients treated with remdesivir are sub-μM (Tempestilli et al., 2020). Indeed, the IC50 of remdesivir against OC43-induced CPE was in the μM range (Fig. 2C-2D). Importantly, co-treatment with either lapatinib or doramapimod reduced the IC50 of remdesivir into the sub-μM range. These results suggest that in addition to blocking OC43 betacoronavirus activity on their own, these compounds could potentially be combined with remdesivir to effectively treat patients infected with SARS-CoV-2.

**Figure 2.**
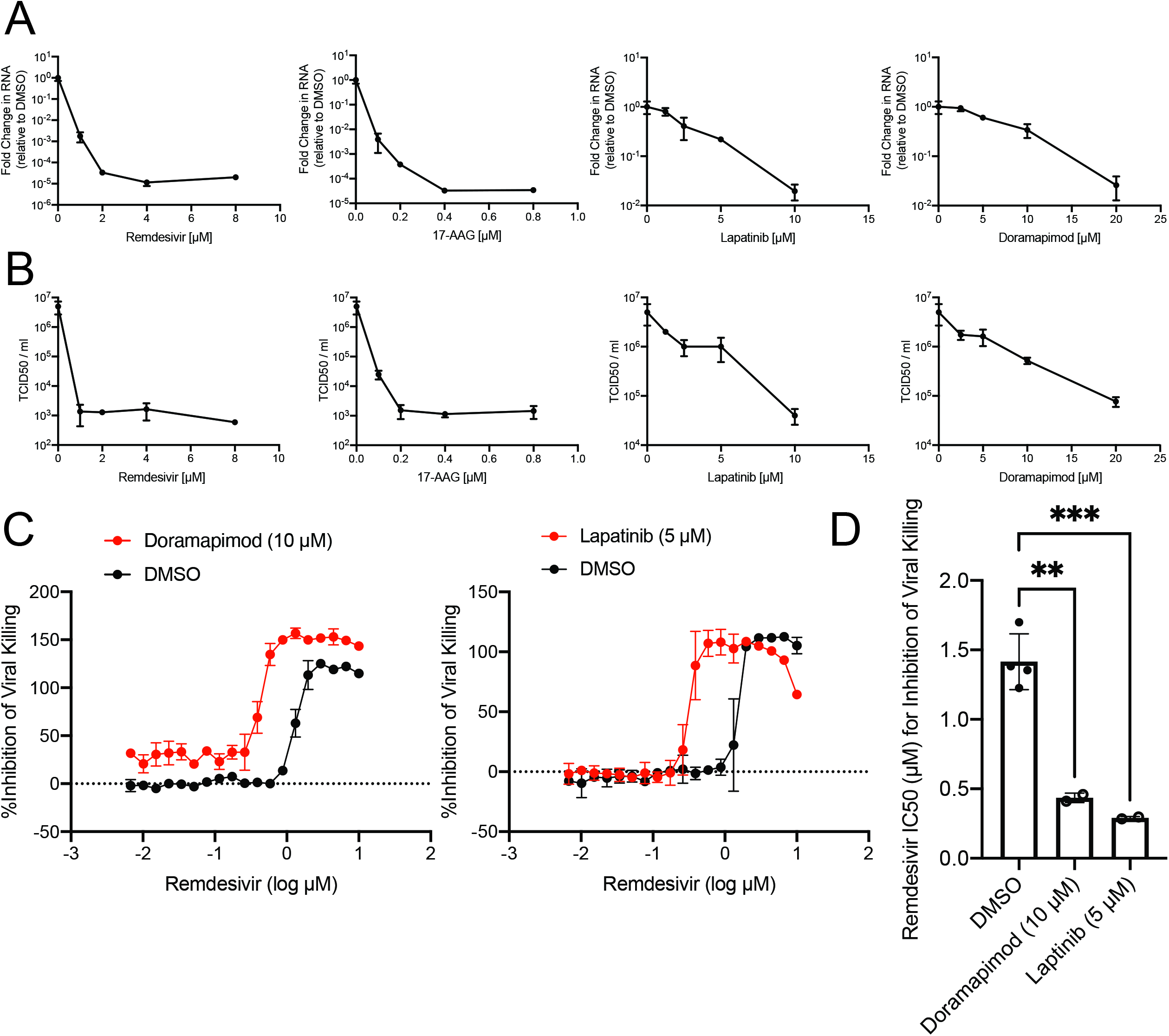
Hit compounds inhibit OC43 replication and synergize with remdesivir. **(A-B)** Confluent MRC5 cells were pretreated with DMSO (0.25%), remdesivir, 17-AAG, lapatinib, or Doramapimod at the indicated concentrations for 3 hours. After pretreatment, cells were infected with OC43 at a MOI of 0.05 in the presence of the appropriate drug concentration. At 24 hpi, RNA **(A)** and virus containing supernatants **(B)** were harvested. cDNA was synthesized from the RNA, and relative OC43 RNA levels were quantified. RNA levels were normalized to levels from DMSO treated samples **(A)**. The virus containing supernatants were s by TCID50 **(B)**. **(C)** Percent inhibition of viral killing was plotted against remdesivir drug concentration either alone (black curve) or in combination with 10 μM doramapimod (left) or 5 μM lapatinib (right). **(D)** IC50 values from **(C)**. \*\**P*<0.01, \*\*\**P*<0.001. Dunnett’s multiple comparisons test was used to determine statistical significance for (**D)**.

### MRC5-ACE2 cells are an effective model of SARS-CoV-2 infection

The betacoronavirus OC43 was used as a surrogate of SARS-CoV-2 for high-throughput screening purposes because it can be safely handled under BSL2 containment (unlike SARS-CoV-2). While our hit compounds displayed impressive antiviral action against OC43, the ultimate goal was to identify therapeutics for SARS-CoV-2 infection. Expression of ACE2 is required for entry of SARS-CoV-2, and overexpression of ACE2 in cells has been effective at permitting infection, although this has largely been examined in transformed cancer cell lines (Dittmar, Jun 19, 2020), which could potentially confound findings. Instead, we decided to express ACE2 in MRC5 cells to create MRC5-ACE2 cells (Fig. 3A). MRC5-ACE2 cells demonstrated robust SARS-CoV-2 infection, as indicated by elevated accumulation of viral RNA, which was comparable to Vero-E6 cells, especially at 24 hpi (Fig. 3B). Further, MRC5- ACE2 cells showed CPE after infection with SARS-CoV-2 at multiple MOIs (Fig. 3C). As MRC5- ACE2 cells are of the appropriate species (human), genetic background (non-transformed), and tissue type (pulmonary), this model represents an important new tool for SARS-CoV-2 research.

**Figure 3.**
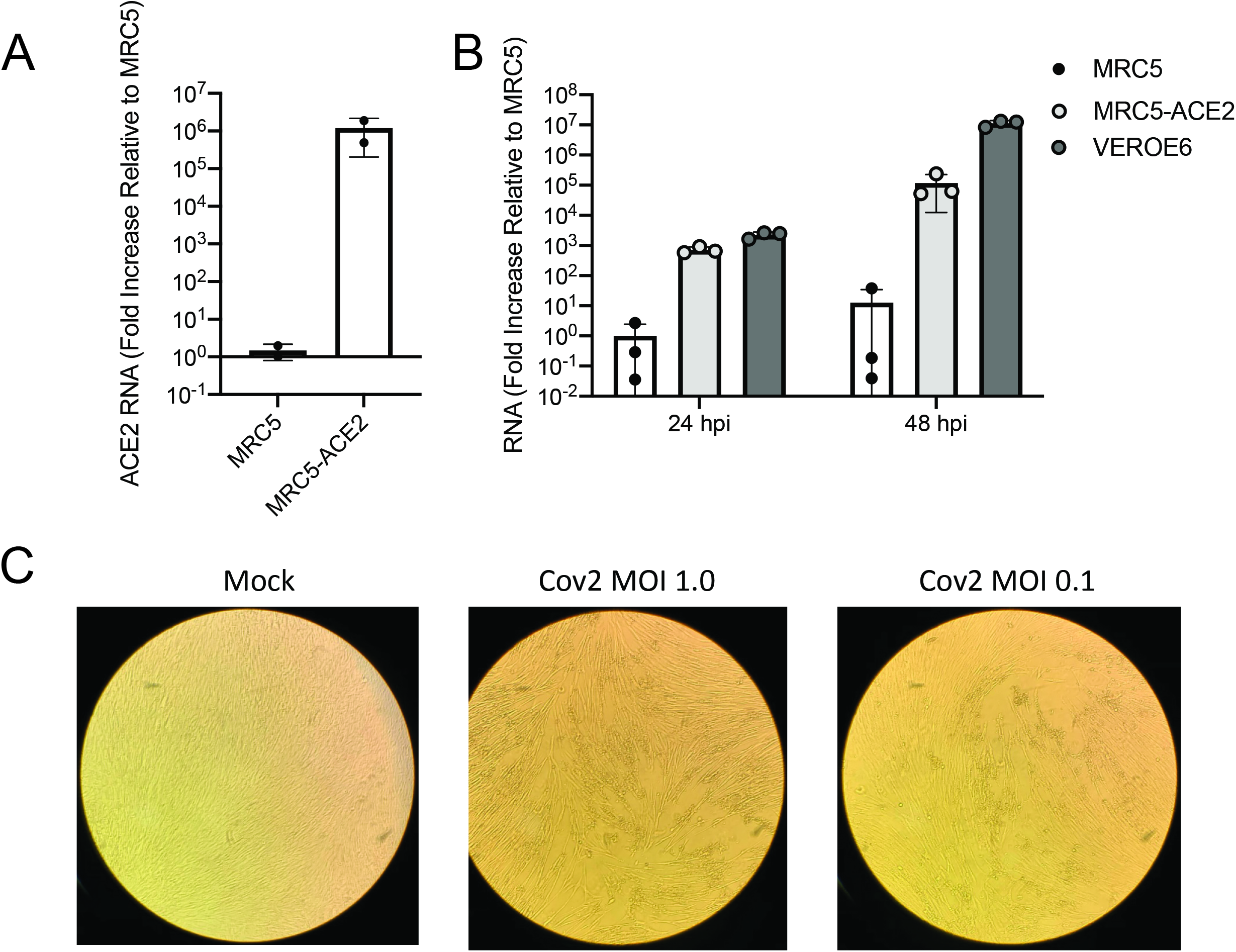
SARS-CoV-2 replicates in MRC5 cells expressing ACE2. **(A)** Confluent MRC5 cells and MRC5 cells expressing ACE2 (MRC5-ACE2) were harvested, and their RNA isolated prior to synthesizing their cDNA. Relative SARS-CoV-2 RNA levels were quantified and normalized to the levels in MRC5 cells. **(B)** Confluent Vero-E6, MRC5, and MRC5-ACE2 cells were infected with SARS-CoV-2 at a MOI of 0.01. At 24 and 48 hpi, RNA was harvested and cDNA was synthesized. Relative SARS-CoV2 RNA levels were quantified and normalized to the levels in MRC5 cells. **(C)** Representative images of MRC5-ACE2 cells either mock infected or infected with SARS-CoV-2 at a MOI of 1.0 or 0.1 at 96 hpi.

### Lapatinib blocks SARS-CoV-2 infection alone or in combination with remdesivir

The hit compounds derived from our MAPS screen with OC43 were lapatinib, doramapimod, and 17-AAG. Since lapatinib and 17-AAG had the most antiviral activity against OC43, these compounds were tested against SARS-CoV-2. Lapatinib effectively blocked SARS-CoV-2 infection, as measured by the dramatic reduction in viral RNA accumulation (Fig. 4A). Interestingly, 17-AAG, which was the top hit against OC43, only minimally reduced SARS-CoV-2 RNA accumulation. There are potentially multiple reasons for the lack of effectiveness observed for 17-AAG against SARS-CoV-2 (dose, timing, differences between betacoronaviruses). Based on the differences in effectiveness of 17-AAG between OC43 and SARS-CoV-2, we decided to examine doramapimod, although it was less effective at blocking OC43 infection. Intriguingly, doramapimod was able to block SARS-CoV-2 infection at a low μM concentration, although it was still not as inhibitory as lapatinib or remdesivir (Fig. 4B). Nonetheless, lapatinib and doramapimod also blocked SARS-CoV-2-induced CPE (Fig. 4C) and completely prevented SARS-CoV-2 N protein accumulation (Fig. 4D). The compounds did not induce any cellular toxicity to MRC5-ACE2 cells at these drug concentrations (Fig. S3). Finally, our hit compounds were combined with remdesivir to examine whether these compounds could be combined to synergistically block SARS-CoV-2 infection. Both lapatinib and doramapimod were able to dramatically reduce the dose required for remdesivir required to abolish SARS-CoV-2 RNA accumulation (Fig. 4E). These findings suggest that our top hit compounds lapatinib and doramapimod, either alone or in combination with remdesivir, are potentially effective therapeutic options for patients infected with SARS-CoV-2.

**Figure 4.**
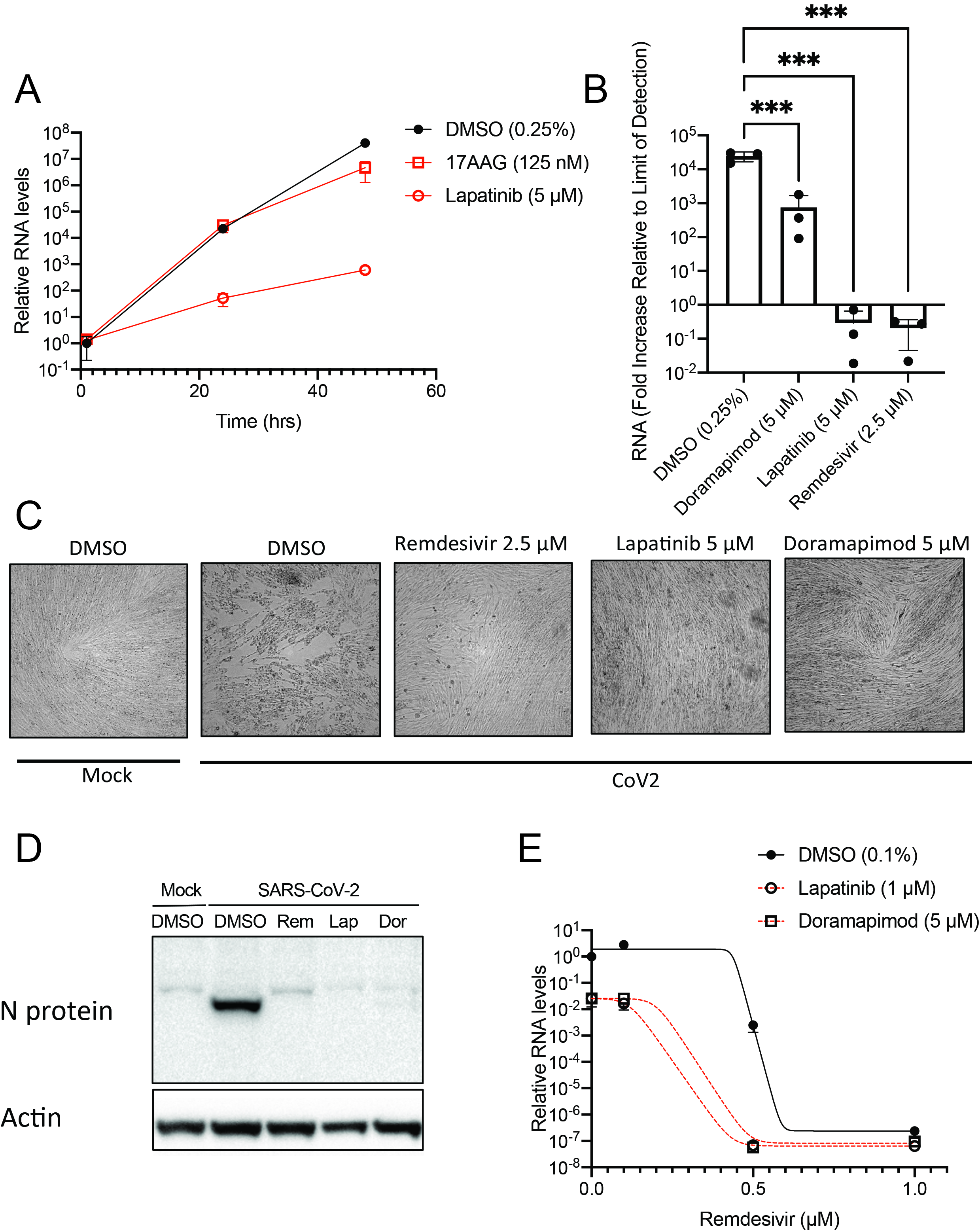
Lapatinib and doramapimod inhibit SARS-CoV-2 replication alone or in combination with remdesivir. **(A)** Confluent MRC5-ACE2 cells were pretreated with DMSO, 17-AAG, or lapatinib at the indicated concentrations for 4 hours. After pretreatment, cells were infected with SARS-CoV-2 at an MOI of 0.01 in the presence of the appropriate drug. At 1, 24, and 48 hpi, RNA was harvested and cDNA was synthesized. Relative SARS-CoV-2 RNA levels were quantified and normalized to levels of the DMSO treated sample at 1 hpi. **(B-C)** Confluent MRC5-ACE2 cells were pretreated with DMSO, doramapimod, remdesivir, or lapatinib at the indicated concentrations for 4 hours. After pretreatment, cells were infected with SARS-CoV-2 at an MOI of 0.01 in the presence of the appropriate drug. At 1 and 48 hpi, RNA was harvested and cDNA was synthesized. Relative SARS-CoV-2 RNA levels at 48 hpi were quantified, and normalized to levels of the DMSO treated sample at 1 hpi **(B)**. At 48 hpi, cells were fixed, and pictures were taken to examine cytopathic effect **(C)**. **(D)** Immunoblot analysis of MRC5-ACE2 cells infected with mock or SARS-CoV-2 (MOI=1) and pretreated with either remdesivir (Rem, 2.5 μM), lapatinib (Lap, 5 μM) or doramapimod (Dor, 5 μM). Cells were harvested at 24 hpi. **(E)** Confluent MRC5-ACE2 cells were pretreated with remdesivir at the indicated concentrations along with DMSO, lapatinib or doramapimod, infected with SARS-CoV-2 at an MOI of 0.01 in the presence of the appropriate drug. At 48 hpi, RNA was harvested and cDNA was synthesized. Relative SARS-CoV-2 RNA levels were quantified and normalized to levels of the DMSO treated sample. \*\*\**P*<<0.001. Dunnett’s multiple comparisons test was used to determine statistical significance for (**B)**. See also **Figure S3**.

## DISCUSSION

Our pharmacological screen identified several compounds capable of blocking the in vitro replication of two betacoronaviruses. We screened against OC43, which is a common human coronavirus that typically causes mild to moderate upper respiratory tract infections. Hit compounds were subsequently found to inhibit SARS-CoV-2, which is responsible for the COVID-19 pandemic and the death of over a million people globally. Of particular interest was the finding that lapatinib could inhibit SARS-CoV-2 replication in pulmonary fibroblasts by over 50,000 fold (Fig 4B). Lapatinib is an FDA-approved compound with a favorable toxicity profile in patients (Baselga et al., 2012). Further, lapatinib concentrations found to inhibit SARS-CoV-2 are readily achievable in human tissues at currently prescribed doses (Spector et al., 2015). These results suggest that lapatinib could be an effective therapeutic to attenuate SARS-CoV-2 associated morbidity and mortality that could be quickly transitioned to the clinic. The need for effective therapeutics to treat SARS-CoV-2 is immense given its devastating effect on global human health, as well the recent disappointing results from the SOLIDARITY trial indicating that other potential antivirals such as remdesivir and hydroxycholroiquine are not clinically effective (Pan et al., 2020).

Traditional drug screening libraries contain a wide range of compounds at a single high concentration dose. While useful for drug discovery with biochemical assays, these libraries are less suited for more complicated screens with live cells in which success depends on multiple parameters that can be affected by drug concentration, e.g. those affecting viral replication or cellular enzymes important for viability. Our current screen examined compounds across a wide range of doses (20 μM to 1 nM), which enabled us to identify concentrations of compounds that inhibit virally-induced killing, as well as those concentrations that contribute to toxicity during mock infection - allowing us to identify a preclinical therapeutic window. Further, traditional screening at a single high dose will miss compounds that are effective antivirals at lower concentrations, but toxic at higher concentrations, i.e., single dose screening increases false negatives. Finally, dose-response screening enables assessment of how partial inhibition of essential activities might impact viral infection, which is a significant limitation of most genetic screening studies (Birsoy et al., 2015; Shalem et al., 2014).

Our results indicate that the hit compounds identified via screening against a minimally pathogenic BSL2 coronavirus can identify compounds that inhibit the more pathogenic SARS-CoV-2 virus, a BSL3 agent. This has significant implications for antiviral development more broadly. Specifically, high-throughput screening at BSL3 or BSL4 conditions is a major impediment to the ability to conduct antiviral screening and thus represents a major barrier to the development of antiviral compounds that could block the most pathogenic viruses. The ability to screen against related BSL2 agents to identify promising compounds could spur the development of compounds against extremely pathogenic agents that require the strictest biocontainment levels.

Many pharmacological screens for SARS-CoV-2 inhibitors have been performed in Vero cells (Bojkova et al., 2020; Gordon et al., 2020), which are monkey kidney cells. Vero cells have many benefits as they are easy to culture, grow quickly, and grow SARS-CoV-2 to high titers, which has been an issue for many human cell lines due to the lack of expression of the SARS-CoV-2 receptor, ACE2. However, screening for antiviral compounds in Vero cells also has limitations. They are derived from an African green monkey *(Cercopithecus aethiops),* from a non-respiratory tissue, i.e., the kidney, and are capable of forming tumors. All of these factors can significantly impact both the cellular and the viral responses to compound treatment. Consistent with this, it has recently been found that many compounds capable of limiting SARS-CoV-2 in Vero cells, such as hydroxychloroquine, are ineffective in human lung epithelial cell lines (Dittmar, Jun 19, 2020). We, therefore, set out to develop a model using human, nontransformed, pulmonary cells. MRC5 cells fulfill these requirements, and after transduction with ACE2 are capable of robust SARS-CoV-2 infection (Fig. 3). We found MRC5 cells to be extremely amenable to high-throughput screening, and given their physiological relevance, i.e., human and non-transformed, they provide a robust platform for future antiviral screening.

While we find that lapatinib attenuates SARS-CoV-2 infection, the mechanisms through which it does so are less clear. Lapatinib is an EGFR/HER2 inhibitor used to treat HER2-positive breast cancer (Burris et al., 2005; Dhillon and Wagstaff, 2007). It is possible that SARS-COV-2 depends on EGFR/HER2 activity or a closely related protein kinase. If such is the case, we might have expected to identify other EGFR-related kinase inhibitors in our screen, e.g., gefitinib and erlotinib. These compounds were not identified, although we cannot rule significant differences in the kinase specificities of the various inhibitors in our library. Another possibility is that lapatinib is inhibiting an alternative target from the one it was designed to block. An in silico molecular docking study identified lapatinib as a putative inhibitor of the SARS-CoV-2 protease, 3CLpro (Ghahremanpour et al., 2020), which is important for productive SARS-CoV-2 infection, and a target for anti-coronaviral therapeutic development (De Clercq, 2006). However, despite these computational predictions, experimental results reported in the same publication suggested that lapatinib did not inhibit 3CLpro (Ghahremanpour et al., 2020). In contrast, a separate manuscript in pre-print suggests that lapatinib could, in fact, inhibit the activity of 3CLpro (Drayman et al., 2020). Regardless of the exact antiviral mechanism, given the immediate need for clinical therapeutics to treat SARS-CoV-2, lapatinib is a good candidate for rapid assessment in Covid-19 patients. Lapatinib is orally bioavailable, and has been found to accumulate in the plasma of human patients at ~1-3 μM concentrations, and achieve ~8-12 μM concentrations in tissue samples (Spector et al., 2015). We observe significant inhibition at these concentrations in vitro suggesting that lapatinib could be effective at concentrations that occur in human patients. Further, lapatinib is well tolerated in patients with mostly mild adverse effects (Dhillon and Wagstaff, 2007). Collectively, the pharmacokinetic and safety profiles of lapatinib suggest that it could be relatively quickly evaluated in the clinic with respect to its ability to attenuate SARS-CoV-2-associated morbidity.

With respect to treatment of SARS-CoV-2, remdesivir, although very effective in vitro, has not been effective in patients (Pan et al., 2020), for uncertain reasons. Remdesivir is a prodrug, that gets transformed into a bioactive nucleoside monophosphate derivative, GS-441524, which targets viral RNA-dependent RNA polymerases. Previous studies have shown that remdesivir can maximally accumulate to approximately 4-9 μM in plasma of healthy patients, while GS-441524 maximally accumulates to concentrations of about 0.5 μM (Jorgensen et al., 2020). In vitro, remdesivir exhibited an EC50 between 0.6-1.5 μM, and GS-441524 demonstrated an EC50 between 0.5-1.1 μM, with different activities in different cell types as a potential explanation for the range of values (Pruijssers et al., 2020). These data suggest that while remdesivir could inhibit SARS-CoV-2 replication at achievable plasma concentrations, the accumulation of its active derivative GS-441524 is likely sub-optimal for its antiviral activity. Further, in remdesivir treated COVID-19 patients, remdesivir was not detectable in bronchoalveolar aspirates, and GS-441524 was only detected at ~15 nM concentrations(Pan et al., 2020), significantly below its EC50. These data suggest that the availability of these compounds to the lung may limit their effectiveness in COVID-19 patients, which could be responsible for their lack of clinical efficacy (Pan et al., 2020). Combinations of antiviral compounds have been very clinically successful in limiting viral pathogenesis, e.g., with HIV-1. Here, we find that both lapatinib and doramapimod are capable of acting synergistically with remdesivir to attenuate SARS-CoV-2 replication, i.e., reducing the concentrations required for their antiviral activity. These results raise the possibility that treatment with lapatinib or doramapimod in combination with remdesivir could substantially improve clinical efficacy. While additional investigations are required, including in vivo validation of drug efficacies, our findings point to novel SARS-CoV-2 treatment options for patients, and potentially improved outcomes during this unprecedented pandemic.

## ACKNOWLEDGMENTS

We would like to thank John Ashton and Tim Bushnell for their support regarding the frequent use of institutional high-throughput screening equipment. The following reagent was obtained through BEI Resources, NIAID, NIH: SARS-Related Coronavirus 2, Isolate Hong Kong/VM20001061/2020, NR-52282. The work was supported by NIH grants AI127370 and AI50698 to J.M and by the American Association for Cancer Research and Breast Cancer Research Foundation (20-20-26-HARR) and Breast Cancer Coalition of Rochester to I.S.H.

## AUTHOR CONTRIBUTIONS

M.H.R., J.H.C., M.M., J.M., and I.S.H. initiated the study and conceived the project, designed the experiments, interpreted the results and wrote the manuscript. M.H.R., J.H.C., and M.M. were involved in conducting all experiments. M.M.L. IV and T.T. assisted with OC43 experiments. L.A.S-S. assisted with high-throughput screening experiments. X.L.S. cloned ACE2 vector and created MRC5-ACE2 cells. J. L., G.A., and S.D. assisted with BSL3 SARS-CoV-2 experiments.

## STAR METHODS

### EXPERIMENTAL MODEL AND SUBJECT DETAILS

#### Cell Culture, Viruses, and Viral Infection

Telomerase-immortalized MRC5 fibroblasts (MRC5 cells) were cultured in Dulbecco’s modified Eagle serum (DMEM; Invitrogen) supplemented with 10% (vol/vol) fetal bovine serum (FBS) (Atlanta Biologicals), 4.5 g/liter glucose, and 1% penicillin-streptomycin (Pen-Strep; Life Technologies) at 37°C in a 5% (vol/vol) CO_2_ atmosphere. Vero-E6 cells were cultured in Eagle’s Minimum Essential Medium (ATCC, #30-2003) supplemented with 10% (vol/vol) fetal bovine serum (FBS) (Atlanta Biologicals), 4.5 g/liter glucose, and 1% penicillin-streptomycin (Pen-Strep; Life Technologies) at 37°C in a 5% (vol/vol) CO_2_ atmosphere. Viral stocks of OC43 were propagated in MRC5 cells in 2% (vol/vol) FBS, 4.5 g/liter glucose, and 1% penicillinstreptomycin at 34 °C. Viral stock titers were determined by TCID50 analysis, and for the assessment of OC43 viral replication, viral titer was determined via TCID50 analysis. The SARS-CoV-2 isolate, Hong Kong/VM20001061/2020, was previously isolated from a nasopharyngeal aspirate and throat swab from an adult male patient in Hong Kong and was obtained through BEI resources (NR-52282). Viral stocks of SARS-CoV-2 were propagated in Vero-E6 cells in 2% (vol/vol) FBS, 4.5 g/liter glucose, and 1% penicillin-streptomycin at 37 °C. Viral stock titers were determined by TCID50 analysis. All experiments involving live SARS-CoV-2 were conducted in a biosafety level 3 facility at the University of Rochester. ACE2-expressing MRC5 cells were generated via lentiviral transduction. For the generation of lentivirus, 293T cells (~50% confluent) were transfected with 2.6 μg pLenti-ACE2, 2.4 μg PAX2, and 0.25 μg vesicular stomatitis virus G glycoprotein using the Fugene 6 transfection reagent (Promega) according to the manufacturer’s protocol. Twenty-four hours later, the medium was replaced with 5 ml of fresh medium. Lentivirus-containing medium was collected after an additional 24 h and filtered through a 0.45μm pore-size filter prior to transduction. Telomerase-immortalized MRC5 fibroblasts were transduced with lentivirus in the presence of 5 μg/ml Polybrene (Millipore Sigma) and incubated overnight. The lentivirus-containing medium was then removed and replaced with fresh medium. The cells were subsequently passaged in the presence of puromycin at 1 ug/ml and remained under selection for 3 days.

### METHOD DETAILS

#### Compound Library and High-Throughput Compound Screening

The compound library for the high-throughput screen was prepared as described (Harris et al., 2019). Prior to screening with the compound library, the robustness of our screening method was determined by obtaining a Z’ factor (Zhang et al., 1999). Each well of a 384-well pate (Corning, 3764) was seeded with 6,000 MRC5 cells in a volume of 30 μL using a Multidrop Combi reagent dispenser. Cells were then incubated at 37°C for 24 hours at which point cells were either given 20 μL of media for a mock infection or 20 μL media containing OC43 to achieve an MOI of 0.05 TCID_50_/mL. Cells were then incubated at 34°C for 96 hours whereupon the average and standard deviation of the number of cells was used to determine the Z’ factor of 0.66. To determine compounds that inhibit the cytopathic effect observed with OC43 infection, 30 μL of MRC5 were seeded per well of 384-well plate and incubated at 37°C. After 24 hours, 100 nL of compounds from the library plates were pin transferred onto the cells and incubated at 34°C for 4 hours at which point 20 μL of media containing OC43 was added to infect the cells at an MOI of 0.05 TCID50/mL. The infected plates were then incubated at 34°C for 96 hours and then cell numbers were quantified. Percent inhibition of viral killing was determined as: (Cell

Number Infected_(Drug)_ - Average Cell Number Infected_(DMSO)_/(Average Cell Number Mock Infection_(DMSO)_ - Average Cell Number Infected_(DMSO)_)^*^100%. All values calculated to be negative were set to “0”. Data post-processing was conducted using R and Prism scripts.

#### Quantification of Cell Numbers

For cell culture experiments in 96-well (Greiner Bio-One, #655160) or 384-well (Corning, #3764) plate formats, cells were washed with PBS (Thermo Fisher, BP399-20), fixed using 4% formaldehyde (Sigma, #252549) and stained with 5 μg/mL Hoechst 33342 (Thermo Fisher). Cells were then imaged using a Cytation 5 imaging reader (BioTek). Each well was imaged using a 4X magnification objective lens and predefined DAPI channel with an excitation wavelength of 377 nm and emission wavelength of 447 nm. Gen5 software (BioTek) was used to determine cell number by gating for objects with a minimum intensity of 3000, a size greater than 5 μm and smaller than 100 μm.

#### Drug Synergy

To identify synergistic effects between the compounds identified in our screen and remdesivir, MRC5 were seeded in a 384-well plate as described for screening and incubated at 37°C for 24 hours. Using a D300 liquid dispenser (Hewlett-Packard), remdesivir was added following 1:2 dilution curve going down the rows of each plate starting at 10 μM in row C and ending at 0.01 μM in row N. Hit compounds were then added using a 19-point dose response curve across the columns with 1:1.5 dilution so that the concentration ranged from 20 μM in column 3 to 0.01 μM in column 21. After compounds were added to the cells, the plates were incubated at 34°C for 4 hours. Plates were then infected with OC43 at an MOI of 0.05 TCID_50_/mL and incubated at 34°C for 96 hours at which point cell numbers were counted.

#### ACE2 Cloning

The human angiotensin-converting enzyme 2 (hACE2) cDNA was amplified by PCR from the hACE2 plasmid (Addgene #1786) using the following primers:

1. 5′-CTT TAA AGG AAC CAA TTC AGT CGA CTG GAT CAT GTC AAG CTC TTC CTG GCT CCT TCT CAG-3′
2. 5′-ACC ACT TTG TAC AAG AAA GCT GGG TCT AGT TAA GCG GGC GCC ACC TGG GAG GTC TCG GTA-3′

PCR-amplified hACE2 cDNA was cloned via Gibson assembly into the BamHI and XbaI sites of pLenti CMV/TO-RasV12-puro backbone (Addgene #22262), to generate pLenti-ACE2.

#### Analysis of Protein Accumulation during SARS-CoV-2 Infections and Drug Treatment

MRC5-ACE2 fibroblast were grown to confluence on a 6-well plate (Greiner Bio-One, #657160) and then pretreated with media containing either DMSO (0.25% Vol/Vol), lapatinib (5 μM), doramapimod (5 μM), or remdesivir (2.5 μM). After 4 hours, cells were either mock infected or infected with SARS-CoV-2, Isolate Hong Kong/VM20001061/2020 (BEI Resources NR-52282) at a MOI of 0.01. After a 1 hour adsorption period, viral inoculum was removed and replaced with media containing drug or DMSO. Samples were harvested in SDS-lysis buffer at 4 and 24 hours post infection. Protein lysates were treated with Laemmli SDS sample buffer (6X; Boston BioProducts, #BP-111R) with 5% β-mercaptoethanol for 10 min at 100°C and then pelleted via centrifugation at 14,500 RPM for 5 minutes. Samples were run on 4-20% polyacrylamide gels (Genescript, #M42015) and transferred onto Immobilon-P PVDF membranes (MilliporeSigma, #ISEQ00010). Membranes were blocked with 5% milk in TBST for 1 hour and blotted overnight with indicated antibodies in TBST with 0.05% bovine serum albumin (BSA). Excess primary antibodies were rinsed away with TBST and rinsed with secondary antibodies in TBST and 0.05% BSA for 1 hour and rinsed again in TBST. Signal was visualized using Clarity Western ECL Substrate (Bio-Rad, #1705060). The following antibodies were used for immunoblot analysis: N-Protein Antibody (SinoBiological, #40068-RP02) and β-Actin (Sigma, #A1978).

#### Analysis of RNA

Before infection with OC43, MRC5 cells were grown to confluence, ~1.2 × 10^5^ cells per cm^2^ in 12-well dishes. Once confluent, cells were pretreated with 400 μl of medium containing drug or DMSO. After three hours, 100 μl of inoculum containing the appropriate drug was added to each well at a multiplicity of infection (MOI) of 0.05. After a 1.5 hour adsorption, 500 ul of media containing the appropriate drug was added to each well for a final volume of 1 mL/well. For infection with SARS-CoV-2, confluent MRC5 cells were pretreated with 400 μl of media or media containing drug or DMSO for four hours. Media was then aspirated and replaced with 200 μl of inoculum containing the appropriated drug at an MOI of 0.01. After 1 hour of adsorption, the inoculum was aspirated and replaced with 1 mL of media or media containing drug or DMSO. OC43 infections were cultured in media containing 10% FBS at 34 °C, while SARS-CoV-2 infections were cultured in media containing 2% FBS at 37 °C. At the indicated times RNA was isolated from infected cells via TRIzol (Invitrogen) extraction, according to the manufacturer’s instructions and used to synthesize cDNA with the qScript cDNA Synthesis Kit (Quantabio). Relative quantities of gene expression were measured and normalized to GAPDH levels via the 2^−ΔΔCT^ method using the following primers: OC43: 5′- GGATTGTCGCCGACTTCTTA-3′ (forward) and 5′-CACACTTCTACGCCGAAACA-3′ (reverse), SARS-CoV-2: 5′-ATGAGCTTAGTCCTGTTG-3′ (forward) and 5′-CTCCCTTTGTTGTGTTGT-3′ (reverse), and human GAPDH: 5′-CATGTTCGTCATGGGTGTGAACCA-3′ (forward) and 5′- ATGGCATGGACTGTGGTCATGAGT-3′. For the analysis of ACE2 expression, total cellular RNA was extracted with TRIzol (Invitrogen) and used to generate cDNA using SuperScript II reverse transcriptase (Invitrogen) according to the manufacturer’s instructions. Transcript abundance was measured by qPCR using Fast SYBR Green master mix (Applied Biosystems), a model 7500 Fast real-time PCR system (Applied Biosystems), and the Fast 7500 software (Applied Biosystems). Gene expression equivalent values were determined using the 2-ΔΔCt method and normalized to GAPDH levels. Biological replicates were analyzed in technical duplicate. Outlier qPCR samples were identified via analysis of GAPDH Ct values that were beyond 1.5 times the interquartile range. For reactions in which the Ct value was undetermined, the limit of detection was set to either a Ct value of 40, or the amount of input inoculum (Ct value of ~37), if it was measured for a given experiment. The expression of hACE2 was confirmed by qPCR using the following primers:

forward primer 5′- GGA GTT GTG ATG GGA GTG ATA G -3′
reverse primer 5′- ATC GAT GGA GGC ATA AGG ATT T -3′

#### Drug Toxicity

MRC5 cells were grown to confluence ~1.2 × 10^5^ cells per cm^2^ in a 12-well dish in media containing 2% FBS at 37 °C. An eight-point 1:2 dilution scheme was made for each drug in media containing 2% FBS using 10 mM stock of drug that was stored in DMSO, starting with the following high concentrations: 17-AAG (1 μM), lapatanib (20 μM), remdesivir (20 μM), doramapimod (20 μM) alongside a 0.25% DMSO (v/v) control. At t0, wells were treated with 1 mL drug of each dilution (n=3). One plate (n=12) was fixed at t0 for a baseline cell count. At t = 48 hpt and t = 96 hpt drug-treated wells were fixed, stained and imaged. Media was aspirated from each well and wells were washed with 500 μl PBS then fixed and stained in 500 μl 4% paraformaldehyde/Hoechst (Invitrogen, #H3570) diluted 1:2000 in PBS for 30 minutes at room temperature. Cells were washed with PBS and nuclei were counted using Cytation 5 imaging software.

### QUANTIFICATION AND STATISTICAL ANALYSIS

#### Statistical Analysis

All statistical analyses were completed using GraphPad Prism 9.0. Sigmoidal 4 point non-linear regression used for **Figure 2C, 4E and S3**.

### SUPPLEMENTAL INFORMATION LEGENDS

**Figure S1.**
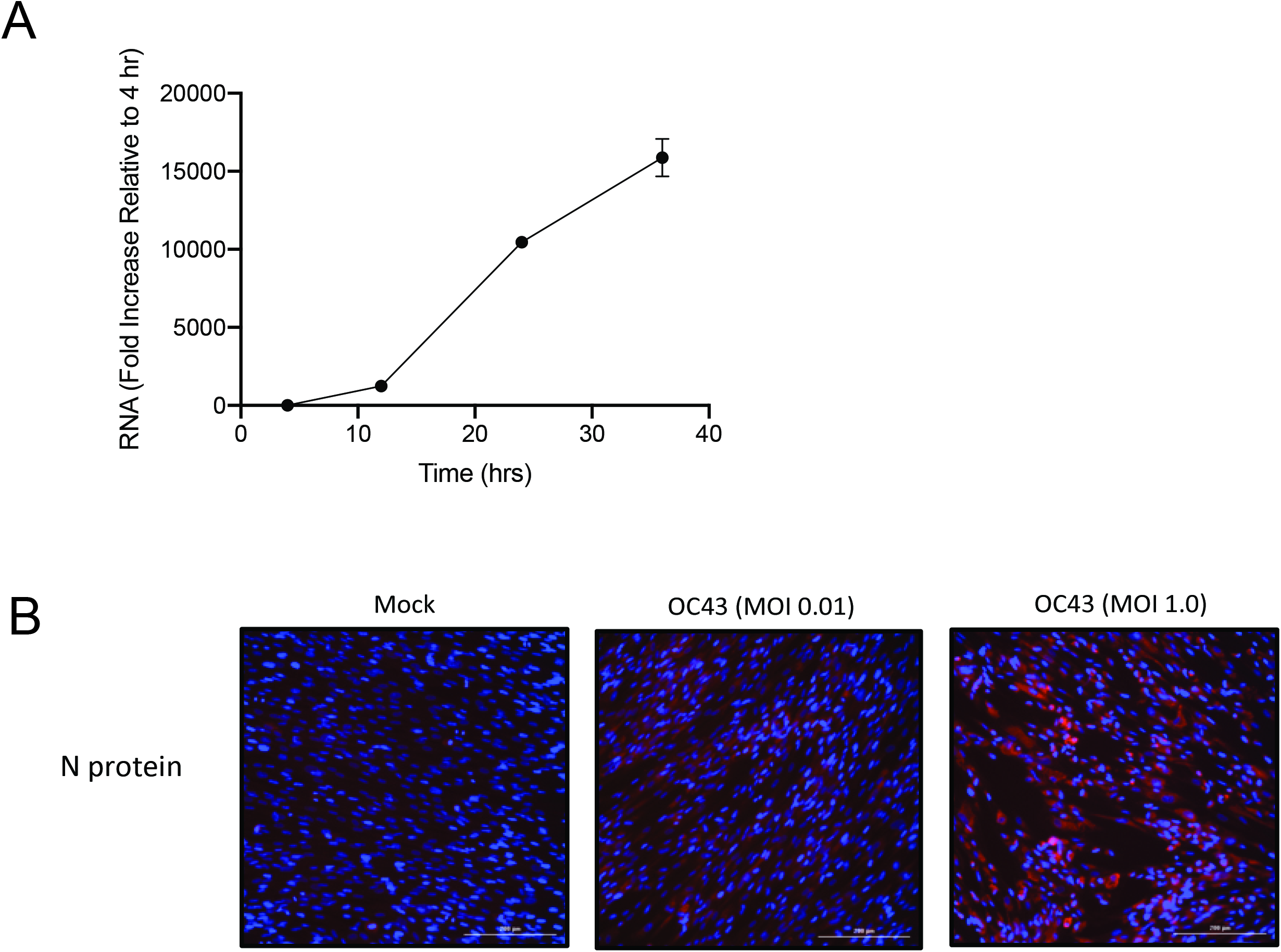
OC43 replicates in MRC5 cells. **(A)**MRC5 cells were infected with OC43 at an MOI of 3.0. At 4, 24, and 72 hpi, RNA was harvested and cDNA was synthesized. Relative OC43 RNA levels were quantified and normalized to levels at 4 hpi. **(B)** MRC5 cells were infected with OC43 at a MOI of 0.01 or 1.0. At 48 hpi, cells were fixed, permeabilized, and immunostained with an antibody specific for N-protein (red). Hoechst fluorescence is shown as blue staining. Scale bars = 200 μm.

**Figure S2.**
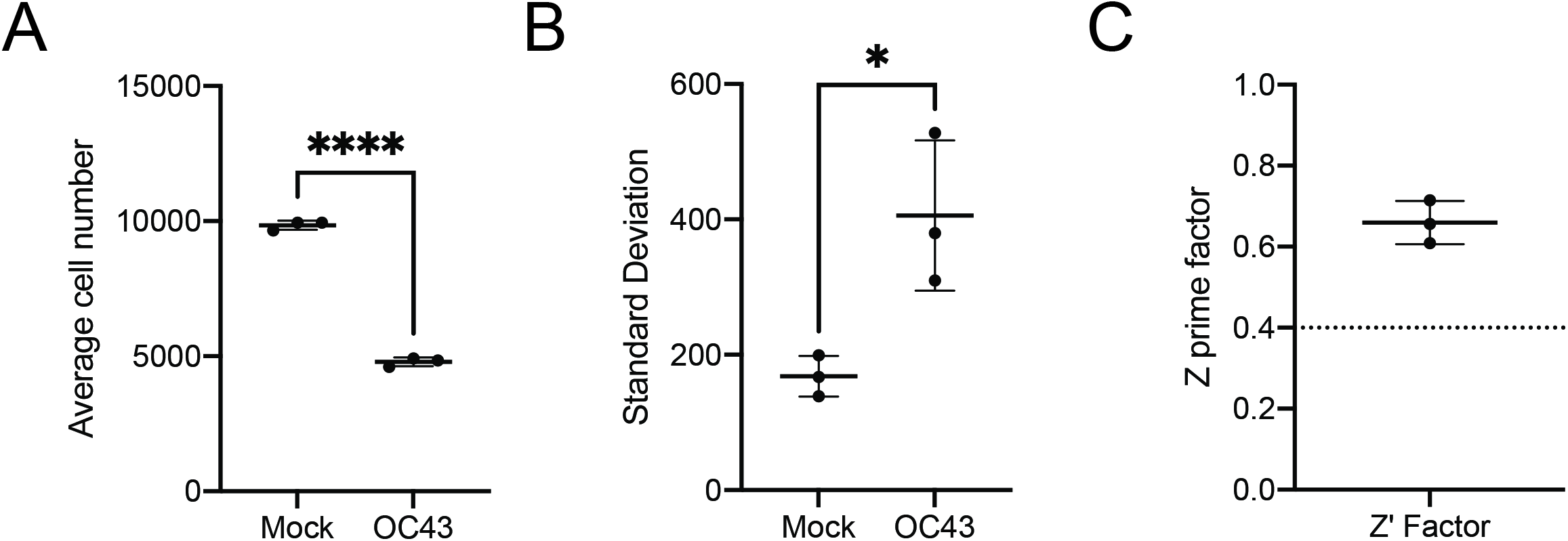
Z prime factor analysis for MRC5 cells with OC43. **(A-C)** Quantification of **(A)** average, **(B)** standard deviation and **(C)** z prime factor from optimization experiments from Figure 1. \**P*<0.05, \*\*\*\**P*<0.0001. Unpaired two-tailed t-test was used to determined statistical significance for **(A-B)**.

**Figure S3.**
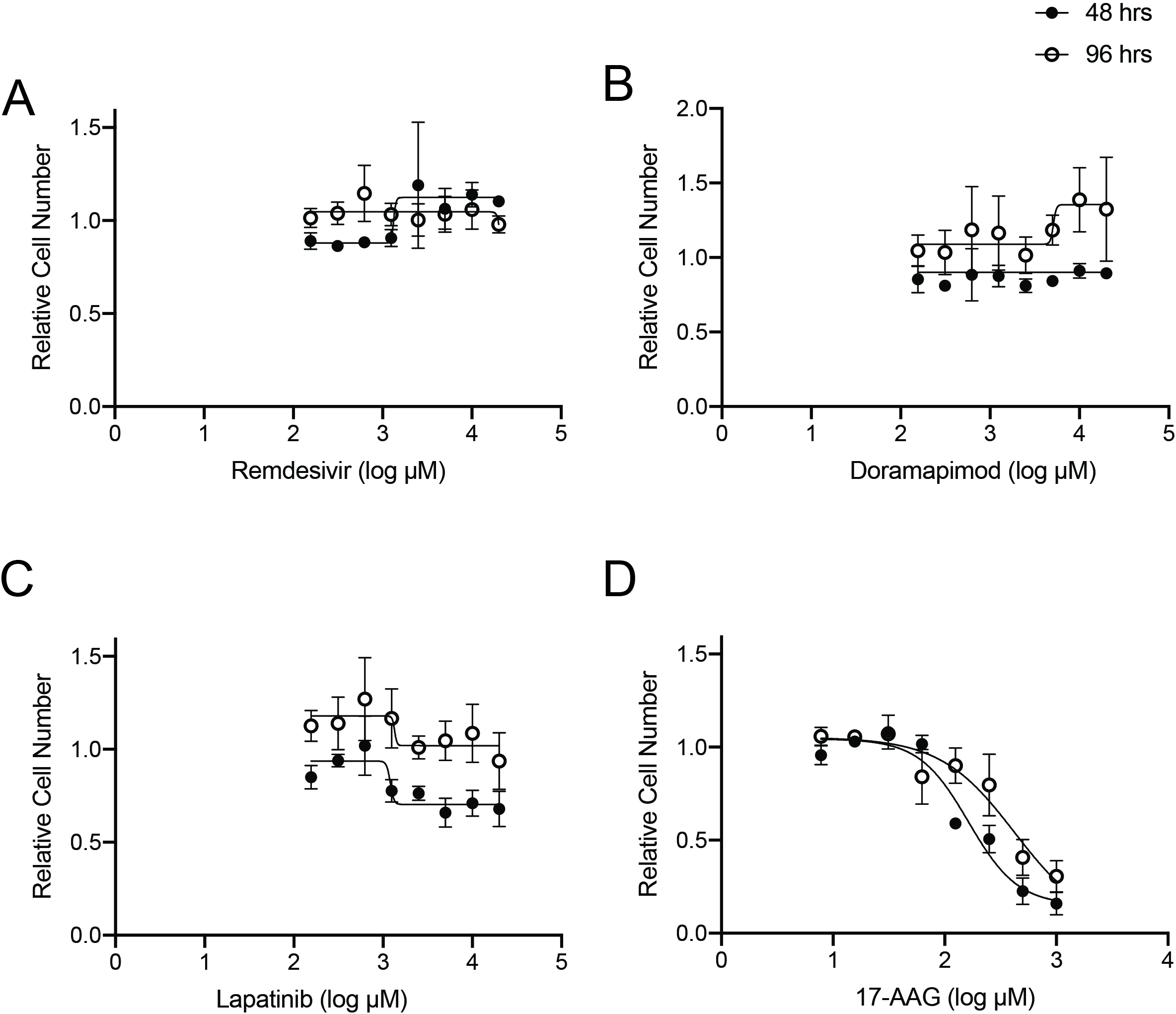
Cytotoxicity of selected drugs in MRC5-ACE2. **(A-D)** MRC5-ACE2 cells were grown to confluence in 12 well dishes and treated with DMSO (0.25%) or the indicated concentration of **(A)** remdesivir, **(B)** doramapimod, **(C)** lapatanib, and **(D)** 17-AAG. At 48 and 96 hours post treatment, cells were fixed in paraformaldehyde and stained with Hoechst. Nuclei were counted for each treatment and relative cell number was normalized to DMSO.

**Table S1. Related to Figure 1.** Maximal percent inhibition of viral killing derived from high-throughput screening of OC43 infection in MRC5 cells.

